# First report of *kdr* L1014F and *ace-1* G119S insecticide resistance in Belgian *Culex* (Diptera: Culicidae) mosquitoes

**DOI:** 10.1101/2022.02.28.482257

**Authors:** Lanjiao Wang, Alina Soto, Laure Remue, Ana Lucia Rosales Rosas, Lander De Coninck, Sam Verwimp, Johanna Bouckaert, Mathias Vanwinkel, Jelle Matthijnssens, Leen Delang

**Affiliations:** Laboratory of Virology and Chemotherapy, Department of Microbiology, Immunology and Transplantation, Rega Institute for Medical Research, KU Leuven, 3000, Leuven, Belgium; Laboratory of Viral Metagenomics, Rega Institute for Medical Research, KU Leuven, 3000, Leuven, Belgium

**Author notes:** Corresponding author: Prof. Leen Delang, Rega Institute for Medical Research, box 1043, Herestraat 49, Leuven 3000, Belgium.

**Keywords:** Insecticide resistance, *kdr* L1014*F*, *ace-1* G119S, multi-resistance, mosquito surveillance

## Abstract

The emergence of West Nile virus and Usutu virus in Europe poses a significant risk to public health. In the absence of efficient antiviral therapy or vaccine candidates, the only strategy to control these arboviruses is to target the *Culex* (Diptera: Culicidae) mosquito vector. However, the selection pressure caused by exposure to insecticides for vector control or agricultural pest control can lead to insecticide resistance, thereby reducing the efficacy of insecticide-based vector control interventions. In *Culex* mosquitoes, two of the most common amino acid substitutions associated with insecticide resistance are the *kdr* L1014F in voltage gated sodium channels and G119S in acetylcholinesterase.

In this study, *Culex pipiens* biotype *pipiens, Culex torrentium*, and *Culex modestus* were sampled from 2019 to 2021 in three distinct environmental habitats (urban, peri-urban, and agricultural) in and around the city of Leuven, Belgium. Individual mosquitoes were screened for two mutations resulting in L1014F and G119S amino acid substitutions. Both mutations were observed in *Culex pipiens* and *Culex modestus* but not in *Culex torrentium* mosquitoes across the four collection sites. Furthermore, multi-resistance or cross-resistance in *Culex pipiens* could be a threat in these areas, as both mutations were observed at low frequencies.

These results provide the first report of *kdr* L1014F and *ace-1* G119S resistance in *Culex pipiens* and *Culex modestus* mosquitoes from Belgium, highlighting the importance of mosquito surveillance to design effective arbovirus outbreak control strategies.

## 1. INTRODUCTION

*Culex* mosquitoes (family: *Culicidae*), such as *Culex pipiens* and *Culex modestus*, are efficient vectors of human and animal pathogens. In Europe, *Culex* mosquitoes are the primary vectors of West Nile virus (WNV) and Usutu virus (USUV), two arthropod-borne viruses (arboviruses) of the *Flaviviridae* family (Fros et al. 2015). Both WNV and USUV are maintained in an enzootic cycle, amplifying in birds and transmitted by mosquitoes to incidental hosts such as humans and horses.

Sporadic outbreaks of WNV in humans have arisen in southern and southwestern Europe since 1963. The most significant outbreak occurred in 2018 with 1,548 locally-acquired cases and 166 deaths reported among several southern European countries (“West Nile virus infection. In: ECDC. Annual epidemiological report for 2018.” 2019). There is increasing evidence that the circulation of WNV is spreading towards northern Europe, and this was exemplified in 2018 in Germany and in 2020 in the Netherlands where the first autochthonous cases of WNV in humans and animals were reported (Bakonyi and Haussig 2020). USUV, on the other hand, has established widespread endemicity in Europe. The virus first emerged in 1996 in Italy and has since spread gradually across the continent causing major outbreaks and mortality in birds (Vilibic-Cavlek et al. 2020). Few human cases of USUV have been reported, yet there is zoonotic potential in areas where competent vector species are present. Furthermore, the co-circulation of WNV and USUV is poorly understood and may threaten public health in the future (Nikolay 2015). A small proportion of WNV and USUV infections in humans results in severe neurological complications and even death, yet there are no efficient antiviral drugs or vaccine candidates available for preventing or treating these virus infections. The emergence of WNV and USUV in Europe thus poses a significant increased risk to public health. However, in the event of an outbreak, the only currently available tools to reduce infections with these viruses are mosquito control interventions.

Adulticiding by fogging is the first-line choice for vector control against *Culex* mosquitoes during emergency situations. This method can rapidly reduce the infectious female mosquito population in treated areas; however, adult control is not recommended for WNV outbreak prevention due to the unpredictable nature of outbreaks (Bellini et al. 2014). Other interventions targeting *Culex* mosquitoes include larval source reduction (such as environmental management and removal of breeding sites), larviciding using chemical or biological agents, and personal protection methods such as repellents and mosquito-proofing homes (Bellini et al. 2014). Pyrethroids, organophosphates, carbamates, and organochlorides are the main chemical classes of insecticides available for use in public health. However, because of long-term safety issues, pyrethroids are the only class allowed for mosquito adulticide in Europe.

A potential consequence of using insecticides for vector control or agricultural pest control is the development of insecticide resistance mechanisms in mosquitoes. The main mechanisms responsible for insecticide resistance in insects include target gene mutations, resulting in amino acid alterations leading to permanent changes in the insecticide target site, increased metabolic activity of enzymes involved in insecticide detoxification, and reduced cuticular penetration of insecticides (Hemingway et al. 2004). Voltage-gated sodium channels allow the passage of sodium ions across the plasma membrane of axons to initiate and propagate action potentials (electrical impulses) which are necessary for controlling insect movement and reaction to stimuli (Dong et al. 2014). When mosquitoes come into contact with pyrethroids and DDT, these molecules bind to sodium channels and prolong their opening, disrupting nerve function and inducing paralysis, knockdown and eventually death. Knockdown resistance (*kdr*) is caused by certain amino acid substitutions in sodium channels, which reduce the sensitivity of these channels to pyrethroids and DDT, thus preventing paralysis and knockdown (Soderlund 2008). The substitution L1014F is a single point mutation at position 1014 in the domain II S6 of sodium channels is most common *kdr* subsititution in *Culex* mosquitoes (Davies et al. 2007). An alternate resistance mechanism is the serine residue of acetylcholinesterase (AChE), the target site for organophosphates and carbamates. In the insect central nervous system, AChE catalyses the hydrolysis of the neurotransmitter acetylcholine to terminate nerve impulses (Hemingway et al. 2004). AChE insensitivity is caused by mutations in the ace-1 gene, resulting in substitutions such as G119S, which reduce the sensitivity of AChE to insecticides in *Culex* and *Anopheles* mosquitoes (Weill et al. 2004).

In Africa, the widespread distribution of long-lasting insecticidal bed nets for malaria control has contributed to the rapid rise and spread of highly insecticide-resistant *Anopheles* mosquitoes (Glunt et al. 2015). As a result, the efficacy of key malaria control interventions was significantly reduced, creating the need for novel combinations of insecticides with differing modes of action to regain the effectiveness of long-lasting insecticidal nets. The widespread impact of insecticide resistance in *Anopheles* mosquitoes highlights the importance of resistance monitoring in all mosquito vectors. The need for increased monitoring and surveillance of European mosquitoes was demonstrated recently in 2016 when pyrethroid resistance was detected in *Aedes albopictus* from Italy, threatening the effectiveness of outbreak response interventions (Pichler et al. 2018). Furthermore, an additional cause for concern is the potential impact of resistance mutations on the efficiency of pathogen transmission (i.e. vector competence). In *Culex quinquefasciatus* mosquitoes it was shown that the presence of one homozygous resistance mutation significantly increased WNV dissemination in the mosquito body, resulting in a higher transmission efficiency than susceptible controls (Atyame et al. 2019). This phenomenon is poorly understood, and the impact of insecticide adaptations on the vector competence for arboviruses has so far only been investigated for three arboviruses including WNV, Zika virus and Dengue virus in distinct mosquito species (Atyame et al. 2019, Deng et al. 2021, Parker-Crockett et al. 2021).

In Belgium, native mosquito species include members of the *Culex pipiens* species complex (*Culex pipiens* biotype *pipiens* and *Culex pipiens* biotype *molestus*), *Culex torrentium*, and *Culex modestus* (Boukraa et al. 2015, Wang et al. 2021a). To date, there is no information regarding the presence of insecticide resistance in mosquito species from Belgium or neighbouring countries. In this study, we investigated the presence of two common insecticide resistance mechanisms, *kdr* L1014F and *ace-1* G119S, in *Culex pipiens* biotype *pipiens, Culex torrentium*, and *Culex modestus* mosquitoes from Belgium. Mosquitoes were sampled in three distinct environmental habitats (urban, peri-urban, and agricultural) in and around the city of Leuven from 2019 to 2021. The insecticide resistance mutations reported here will provide insight for evidence-based vector control for the prevention and mitigation of arbovirus outbreaks in Belgium.

## 2. MATERIALS AND METHODS

### 2.1 Ethics statement

Permits for field collections in private habitats (Bertem and Mechelen) were obtained from the landowners. Permits for collection in the Botanic Garden were obtained from the City Green Management of Leuven. Permits for collection in Arenberg Park were obtained from the security responsible of KU Leuven.

### 2.2 Mosquito collections

Collections were performed from August to the beginning of October in 2019, 2020 and 2021, when the weather was adequate, without strong wind or heavy rain. Adult mosquitoes were collected using BG-Sentinel traps (Biogents^®^ AG, Regensburg, Germany) baited with dry ice for CO2 release and BG-lure (Biogents^®^ AG, Regensburg, Germany) to simulate attractive host scent. The traps were emptied and repositioned between sunrise and sunset of the next day. All captured insects were transported to the insectary facility at the Rega Institute, KU Leuven. Mosquitoes were anesthetized and sorted on dry ice and then separated to genus level based on morphological characteristics. Mosquitoes were stored individually at -80°C until further processing. Ten traps were rotated between three collection sites in Leuven and one site in Mechelen (approximately 30 km from Leuven): Leuven urban habitat (Botanic Garden of Leuven, N 50°52’41, E 4°41’21), Leuven peri-urban habitat (Arenberg Park of KU Leuven, N 50°51’46, E 4°41’01), Leuven agricultural habitat (Bertem, N 50°51’57, E 4°37’53), and Mechelen peri-urban habitat (N 51°02’34, E 4°29’08).

### 2.3 DNA extraction & mosquito identification

#### DNA extraction

Following morphological identification, mosquitoes were transferred to tubes with 2.8 mm ceramic beads (Precellys^®^, Bertin, USA) and 500 μl phosphate-buffered saline (1X PBS) solution and homogenised with a Precellys Evolution homogeniser at 2 cycles of 6800 rpm for 10 sec with a pause of 20 sec. The homogenate was lysed at 100°C for 10 min, followed by centrifugation at 10,000 rpm for 1 min to spin down the tissue debris, and 150 μl of supernatant was transferred to a new tube. DNA extraction was performed using the QIAamp DNA kit (Qiagen^®^, Hilden, Germany) according to the manufacturer’s protocol.

#### Mosquito identification

*Culex pipiens* biotype *pipiens* and *Culex pipiens* biotype *molestus* were distinguished with a multiplex qRT-PCR using primers and probes described previously (Rudolf et al. 2013). The biotyping qRT-PCR was confirmed on a subset of samples using single sequencing of the mosquito cytochrome oxidase 1 (COX1) gene. A 25 μl reaction volume was prepared for each reaction using the Low ROX One-Step qRT-PCR 2X MasterMix kit (Eurogentec^®^, Seraing, Belgium) following the manufacturer’s instructions. The cycle conditions were as follows: initial denaturation at 95°C for 10 min, 40 cycles of denaturation at 94°C for 40 sec, elongation at 48°C for 1 min, and extension at 72°C for 1 min, and a final hold stage at 72°C for 2 min. *Culex modestus* and *Culex torrentium* mosquitoes were identified morphologically using the key of Becker (Becker et al. 2010) followed by COX1 Sanger sequencing (Wang et al. 2021a).

### 2.4 PCR & gel electrophoresis

#### Detection of kdr L1014F (mutation from TTA to TTT)

The protocol for *kdr* L1014F detection was adapted from a protocol described elsewhere (Martinez-Torres et al. 1999). Briefly, two PCR reactions were run in parallel using 4 primers: Cgd1 (5′-GTGGAACTTCACCGACTTC-3′), Cgd2 (5′ GCAAGGCTAAGAAAAGGTTAAG-3′), Cgd3 (5′-CCACCGTAGTGATAGGAAATTTA-3′) and Cgd4 (5′-CCACCGTAGTGATAGGAAATTTT-3′). A 20 μl reaction volume was prepared for each reaction using the GoTaq^®^ Green Master 2X Mix (M7122, Promega, Belgium) following the manufacturer’s instructions. The first reaction contained the primers Cgd1, Cgd2 and Cgd3 to identify the presence of the L1014 susceptible allele, and the second reaction consisted of Cgd1, Cgd2 and Cgd4 to identify the presence of the F1014 resistant allele. Taken together, the results of both reactions were used to determine the genotype for each individual as susceptible homozygote (L/L: SS), resistant heterozygote (L/F: RS), or resistant homozygote (F/F: RR). The cycle conditions were as follows: initial denaturation at 94°C for 2 min, 40 cycles of denaturation at 94°C for 30 sec, elongation at 48°C for 30 sec, extension at 72°C for 1 min, and a final hold stage at 72°C for 10 min. Amplicons were separated by electrophoresis on 1.5% agarose gel and were visualised by Midori green staining under UV light with FAS-V Imaging System.

#### Detection of ace-1 G119S (mutation from GGC to AGC)

This analysis was performed following a protocol described previously (Weill et al. 2004) with minor modifications. The DNA block was amplified using the primers CxEx3dir (5′-CGACTCGGACCCACTGGT-3′) and CxEx3rev (5′-GTTCTGATCAAACAGCCCCGC-3′) and a 20 μl reaction volume was prepared for each reaction using the GoTaq^®^ Green Master 2X Mix (M7122, Promega, Belgium) following the manufacturer’s instructions. The cycle conditions were as follows: initial denaturation at 94°C for 2 min, 40 cycles of denaturation at 94°C for 30 sec, elongation at 54°C for 30 sec, extension at 72°C for 1 min, with a final hold stage at 72°C for 10 min. The amplicons were digested with AluI restriction enzyme (Jena Science, Sapphire, USA) following the manufacturer’s instructions and separated on a 2% agarose gel. The products were visualised by Midori green staining under UV light with FAS-V Imaging System. The results were used to determine the genotype for each individual as susceptible homozygote (G/G: SS), resistant heterozygote (G/S: RS), or resistant homozygote (S/S: RR).

### 2.5 Statistical analysis

For the distribution of genotypic and allelic frequencies resulting in the *kdr* L1014F or *ace-1* G119S, analysis was performed among different collecting sites, years, and species by GraphPad Prism 9 (V.9.3). A Hardy–Weinberg equilibrium of the resistant (R) and susceptible (S) allelic frequencies was evaluated using the equations below on the data from *Culex pipiens* collected from 2019 to 2021. The *χ*^2^ and *p*-values were calculated with GraphPad Prism (V.9.3). A p-value of <0.05 was considered statistically significant.

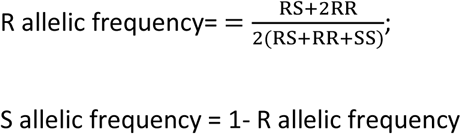

## 3. RESULTS

### 3.1 Frequency and stability of *kdr* over 3 years

*Culex pipiens* were captured from 2019 to 2021 at the urban and peri-urban sites in Leuven. The collection of *Culex pipiens* expanded to the Leuven agricultural site from 2020 to 2021 and to the Mechelen peri-urban site in 2021. *Culex modestus* were collected at the Leuven peri-urban site in 2019, and *Culex torrentium* were captured at the Leuven urban and peri-urban sites in 2020 and the Leuven urban site in 2021. One hybrid female *Culex pipiens* (hybrid of *pipiens* and *molestus*) and one female *Culex pipiens molestus* were identified at the Leuven peri-urban site and Mechelen peri-urban site in 2021, respectively.

The mutation frequencies at position 1014 of the domain II S6 of the sodium channel of *Culex pipiens, Culex modestus* and *Culex torrentium* mosquitoes are presented in Table 1. The most common allele combination for *Culex pipiens* and *Culex modestus* was the susceptible L/L, followed by resistant heterozygosity (L/F). Resistant homozygosity (F/F) presented at a low frequency in *Culex pipiens*, whereas no homozygous resistance genotype was detected in *Culex modestus*, and no resistant heterozygosity or homozygous resistance was detected in *Culex torrentium*.

**Table 1.** Total number and frequency (%) of L/L (susceptible), L/F (heterozygous resistant) or F/F (homozygous resistant) *kdr* alleles in *Culex* mosquitoes collected in the Leuven urban (U), peri-urban (PU), and agricultural (A) sites and the Mechelen peri-urban (MP) site from 2019 to 2021.

| Year | Site | No. | *Culex pipiens* |  |  | No. | *Culex modestus* |  |  | No. | *Culex torrentium* |  |  |
| --- | --- | --- | --- | --- | --- | --- | --- | --- | --- | --- | --- | --- | --- |
|  |  |  | L/L% | L/F% | F/F% |  | L/L% | L/F% | F/F% |  | L/L% | L/F% | F/F% |
|  |  |  | (no.) | (no.) | (no.) |  | (no.) | (no.) | (no.) |  | (no.) | (no.) | (no.) |
| 2019 | PU | 8 | 75 | 12.5 | 12.5 | 51 | 57 | 43 | 0 | - | - | - | - |
|  |  |  | (6) | (1) | (1) |  | (29) | (22) | (0) |  |  |  |  |
|  | U | 12 | 67 | 25 | 8.3 | - | - | - | - | - | - | - | - |
|  |  |  | (8) | (3) | (1) |  |  |  |  |  |  |  |  |
| 2020 | PU | 33 | 73 | 24 | 0.3 | - | - | - | - | 2 | 100 | 0 | 0 |
|  |  |  | (24) | (8) | (1) |  |  |  |  |  | (2) | (0) | (0) |
|  | A | 30 | 77 | 23 | 0 | - | - | - | - | - | - | - | - |
|  |  |  | (23) | (7) | (0) |  |  |  |  |  |  |  |  |
|  | U | 32 | 56 | 34 | 9.4 | - | - | - | - | 4 | 100 | 0 | 0 |
|  |  |  | (18) | (11) | (3) |  |  |  |  |  | (4) | (0) | (0) |
| 2021 | PU | 30 | 87 | 10 | 3.3 | - | - | - | - | - | - | - | - |
|  |  |  | (26) | (3) | (1) |  |  |  |  |  |  |  |  |
|  | A | 30 | 63 | 37 | 0 | - | - | - | - | - | - | - | - |
|  |  |  | (19) | (11) | (0) |  |  |  |  |  |  |  |  |
|  | U | 28 | 64 | 29 | 7.1 | - | - | - | - | 2 | 100 | 0 | 0 |
|  |  |  | (18) | (8) | (2) |  |  |  |  |  | (2) | (0) | (0) |
|  | MP | 33 | 85 | 9.1 | 6.0 | - | - | - | - | - | - | - | - |
|  |  |  | (28) | (3) | (2) |  |  |  |  |  |  |  |  |

The observed frequency of resistance genotypes in *Culex pipiens* was consistent over time in each collection site. The Hardy-Weinberg equilibrium was used to measure the stability of mutations in the population of *Culex pipiens* from all locations over time (Table 2). Ultimately, the difference between the observed and expected *kdr* L1014F genotype frequencies each year were not significantly different and are therefore expected to remain in equilibrium (constant).

**Table 2.** Hardy-Weinberg equilibrium of *kdr* L/L (susceptible), L/F (heterozygous resistant) or F/F (homozygous resistant) alleles in *Culex pipiens*.

| Year | No. observed | | | No. expected | | | $\chi^2$, *p*-value |
| --- | --- | --- | --- | --- | --- | --- | --- |
|  | L/L | L/F | F/F | L/L | L/F | F/F |  |
| 2019 | 14 | 4 | 2 | 12.80 | 6.40 | 0.80 | 2.813, *p*=0.245 |
| 2020 | 65 | 26 | 4 | 64.04 | 27.92 | 3.04 | 0.450, *p*=0.799 |
| 2021 | 91 | 25 | 5 | 88.53 | 29.94 | 2.53 | 3.295, *p*=0.193 |

No. observed: total mosquitoes captured (all locations grouped). No. expected: expected number of mosquitoes based on the Hardy-Weinberg model. The $\chi^2$ and *p*-values were calculated with GraphPad Prism (V.9.3). A *p*-value of <0.05 was considered statistically significant.

### 3.2 Frequency and stability of AChE insensitivity over 3 years

The mutation frequencies in the *ace-1* gene resulting in the G119S amino acid change in *Culex pipiens, Culex modestus* and *Culex torrentium* from 2019-2021 are presented in Table 3. In *Culex pipiens*, AChE susceptibility (G/G) was detected at a higher frequency to heterozygous resistance (G/S) at the urban site (2019-2021) and peri-urban site (2020) in Leuven and the peri-urban site in Mechelen (2021). An equal proportion of susceptible (G/G) and heterozygous resistant (G/S) mosquitoes were observed at the Leuven peri-urban site in 2019 and again in 2021. The rate of heterozygous resistance (G/S) surpassed the rate of full susceptibility (G/G) in *Culex pipiens* at the agricultural site from 2020 to 2021. AChE homozygous resistance (S/S) was only observed in a single mosquito captured at the Mechelen peri-urban site (2021). Almost all captured *Culex modestus* had the full susceptibility (G/G) genotype, yet a small minority presented with heterozygous resistant (G/S) alleles (Leuven peri-urban site, 2019). All captured *Culex torrentium* at the urban (2020-2021) and peri-urban (2020) sites in Leuven were observed with susceptible (G/G) AChE alleles.

**Table 3.** Total number and frequency (%) of G/G (susceptible), G/S (heterozygous resistant) or S/S (homozygous resistant) *AChE* alleles in *Culex* mosquitoes collected in the Leuven urban (U), peri-urban (PU), and agricultural (A) sites and the Mechelen peri-urban (MP) site from 2019 to 2021.

| Year | Site | No. | *Culex pipiens* |  |  |  | *Culex modestus* |  |  |  | *Culex torrentium* |  |  |
| --- | --- | --- | --- | --- | --- | --- | --- | --- | --- | --- | --- | --- | --- |
|  |  |  | G/G% | G/S% | S/S% | No. | G/G% | G/S% | S/S% | No. | G/G% | G/S% | S/S% |
|  |  |  | (no.) | (no.) | (no.) |  | (no.) | (no.) | (no.) |  | (no.) | (no.) | (no.) |
| 2019 | PU | 8 | 50 | 50 | 0 | 51 | 96 | 3.9 | - | - | - | - | - |
|  |  |  | (4) | (4) | (0) |  | (49) | (2) |  |  |  |  |  |
|  | U | 12 | 75 | 25 | 0 | - | - | - | - | - | - | - | - |
|  |  |  | (9) | (3) | (0) |  |  |  |  |  |  |  |  |
| 2020 | PU | 33 | 73 | 27 | 0 | - | - | - | - | 2 | 100 | 0 | 0 |
|  |  |  | (24) | (9) | (0) |  |  |  |  |  | (2) | (0) | (0) |
|  | A | 30 | 47 | 53 | 0 | - | - | - | - | - | - | - | - |
|  |  |  | (14) | (16) | (0) |  |  |  |  |  |  |  |  |
|  | U | 32 | 62.5 | 37.5 | 0 | - | - | - | - | 4 | 100 | 0 | 0 |
|  |  |  | (20) | (12) | (0) |  |  |  |  |  | (4) | (0) | (0) |
| 2021 | PU | 30 | 50 | 50 | 0 | - | - | - | - | - | - | - | - |
|  |  |  | (15) | (15) | (0) |  |  |  |  |  |  |  |  |
|  | A | 30 | 43 | 57 | 0 | - | - | - | - | - | - | - | - |
|  |  |  | (13) | (17) | (0) |  |  |  |  |  |  |  |  |
|  | U | 28 | 75 | 25 | 0 | - | - | - | - | 2 | 100 | 0 | 0 |
|  |  |  | (21) | (7) | (0) |  |  |  |  |  | (2) | (0) | (0) |
|  | MP | 33 | 64 | 33 | **3.0** | - | - | - | - | - | - | - | - |
|  |  |  | (21) | (11) | (1) |  |  |  |  |  |  |  |  |

The observed rate of heterozygosity (G/S) in *Culex pipiens* was generally consistent over time across all locations, except at the Leuven peri-urban site where the rate of heterozygosity fell from 50% to 27% from 2019 to 2020 but returned to 50% in 2021. The Hardy-Weinberg equilibrium of genotypes in *Culex pipiens* across all collection sites remained constant from 2019 to 2020 (Table 4). However, in 2021 the rates of G/G and S/S genotypes were lower and the rate of G/S heterozygosity was higher than the expected frequencies. Therefore, we may observe a higher rate of *ace-1* G119S heterozygous resistance in *Culex pipiens* in this region in the near future.

**Table 4.** Hardy-Weinberg equilibrium of G/G (homozygous susceptible), G/S (heterozygous resistant) or S/S (homozygous resistant) AChE alleles in *Culex pipiens*.

| Year | No. observed | | | No. expected | | | $\chi^2$, $p$-value |
| --- | --- | --- | --- | --- | --- | --- | --- |
|  | G/G | G/S | S/S | G/G | G/S | S/S |  |
| **2019** | 13 | 7 | 0 | 13.6 | 5.8 | 0.6 | 0.875, $p=0.646$ |
| **2020** | 58 | 37 | 0 | 61.6 | 29.8 | 3.6 | 5.550, $p=0.062$ |
| **2021** | 70 | 50 | 1 | 74.6 | 40.8 | 5.6 | 6.137, $p=0.047^*$ |

No. observed: total mosquitoes captured (all locations grouped). No. expected: expected number of mosquitoes based on the Hardy-Weinberg model. The $\chi^2$ and $p$ -values were calculated with GraphPad Prism (V.9.3). A $p$ -value of $<0.05$ was considered statistically significant, (\* = $p < 0.05$ ).

### 3.3 Distribution of L1014F and G119S in *Culex pipiens* collected from different habitat types

In this study, three habitat types including urban, peri-urban, and agricultural were chosen for mosquito collections. For L1014F *kdr* allele detection, the L/L (susceptible homozygote) combination was most dominant (>60%) in all collection sites. At the Leuven peri-urban site, L/F (resistant heterozygote: 15%) was less frequently observed than at the other two collection sites in Leuven (around 30%), but this difference was not significant. In addition, no F/F (resistant homozygote) was observed in the agricultural habitat (Figure 1, A). For the *ace-1* G119S genotype, the proportion of G/S (resistant heterozygote) alleles was lower in the urban site (25-37.5%) than in the agricultural area (53-57%) and in peri-urban sites (33% -50%) (Figure 1, B), however, more samples were needed to draw a significant conclusion. The only mosquito harbouring the S/S (resistant homozygote) allele was observed in Mechelen. Interestingly, similar proportions of both mutations were observed in the periurban area of Mechelen (Figure 1, A and B).

**Figure 1.**
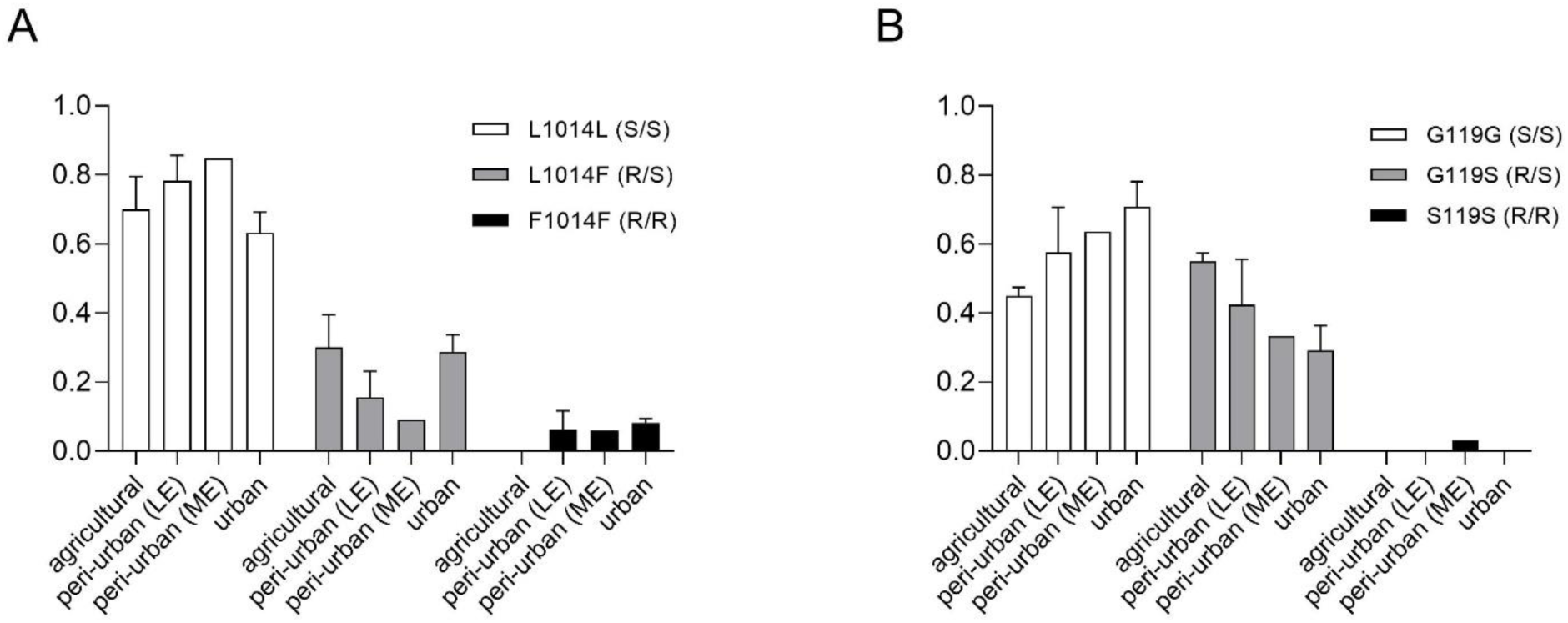
Distribution of *kdr* L1014F and *ace*-1 G119S in mosquitoes from different locations. (A) Prevalence of mosquitoes with L/L (homozygous susceptible; S/S), L/F (heterozygous resistant; R/S) or F/F (homozygous resistant; R/R) *kdr* genotypes. (B) Prevalence of mosquitoes with G/G (homozygous susceptible; S/S), G/S (heterozygous resistant; R/S) or S/S (homozygous resistant; R/R) *AChE* genotypes. Urban (Leuven)= botanic garden, peri-urban =Arenberg Park, agricultural =Bertem. LE= Leuven, ME=Mechelen.

### 3.4 Frequency of L1014F and G119S multi-resistance

When combining the two resistance mechanisms in the *Culex pipiens* sampled in this study, there are seven possible genotypic profiles: L/L-G/G (SSSS), L/L-G/S (SSRS), L/L-S/S (SSRR), L/F-G/G (RSSS), L/F-G/S (RSRS), F/F-G/G (RRSS) and F/F-G/S (RRRS) (Figure 2). The two genotypes RSRR and RRRR were not identified among the sampled Belgian mosquitoes. The SSSS genotype was the most prevalent (>40%) in the collections from the urban and peri-urban sites in Leuven and the peri-urban site in Mechelen. However, from the agricultural site, the resistant genotypes were likely more established, with the SSRS genotype (40%) surpassing the SSSS genotype (23%-36%) in frequency, but not significantly when compared to other locations (Figure 2). In *Culex modestus*, resistant homozygotes were not observed. Only four genotypes were found in *Culex modestus* including SSSS (53%), SSRS (2%), RSSS (43%), and RSRS (2%). In *Culex torrentium*, only the SSSS profile was detected. Interestingly, the *Culex pipiens, Culex modestus* and *Culex torrentium* mosquitoes collected from the same location (Leuven peri-urban site) were found to have different genotypes.

**Figure 2.**
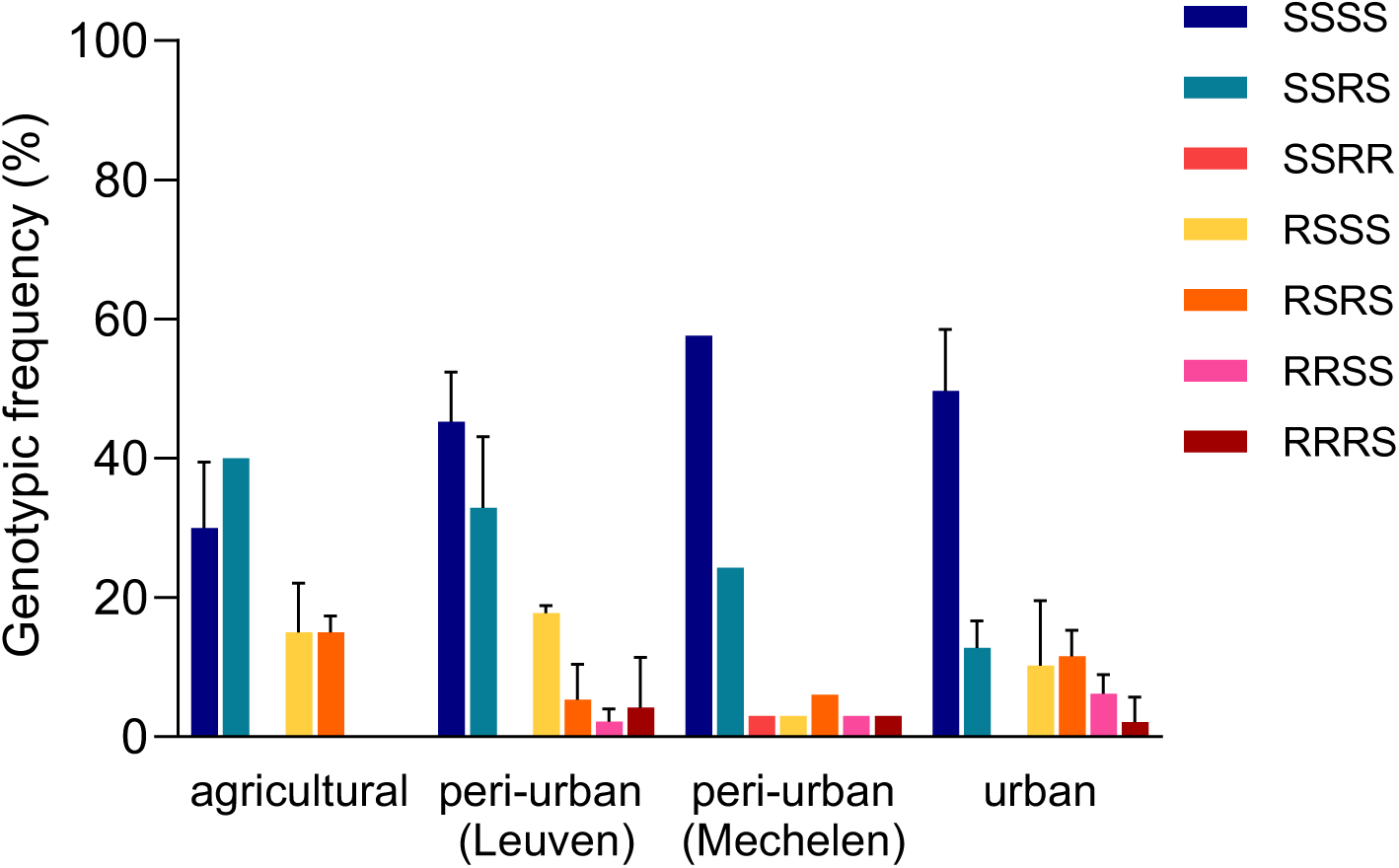
Frequency of combinations of *kdr* L1014F and *ace*-1 G119S in *Culex pipiens* mosquitoes from different locations. A four-letter abbreviation was used to designate the combinations of genotypes: the first two letters refer to *kdr* L1014F and the last two refer to *ace*-1. SS susceptible genotypes, RS heterozygous genotypes, RR homozygous resistant genotypes. L/L-G/G (SSSS), L/L-G/S (SSRS), L/L-S/S (SSRR), L/F-G/G (RSSS), L/F-G/S (RSRS), F/F-G/G (RRSS) and F/F-G/S (RRRS). Urban (Leuven)= botanic garden, peri-urban =Arenberg Park, agricultural =Bertem.

## 4. DISCUSSION

Field mosquitoes were collected in Leuven and surrounding areas in urban, peri-urban, and agricultural habitats during the summer months for three consecutive years (2019-2021). *Culex pipiens* was the dominant species across all collection sites, a finding consistent with previous research conducted in this area (Wang et al. 2021b). The majority of *Culex pipiens* were biotype *pipiens*. Only one hybrid female *Culex pipiens* and one female *Culex pipiens molestus* were identified in this study. *Culex pipiens* biotype *pipiens* and biotype *molestus* can present genetic, behavioural and physiological differences. More importantly, vector competence can vary significantly between the two biotypes; therefore, the identification of the subspecies level is important to assess the risk of arbovirus transmission by field *Culex pipiens* mosquitoes (Abbo et al. 2021).

The *kdr* L1014F and *ace-1* G119S amino acid substitutions were previously observed in *Culex pipiens* from Morocco (Bkhache et al. 2016), Greece (Kioulos et al. 2014), Turkey (Guz et al. 2020), California, USA (Yoshimizu et al. 2020) and China (Liu et al. 2019). *Culex pipiens* sampled from three diverse locations in Greece had a L1014F mutation resistant frequency of 97% (40% heterozygous L/F and 57% homozygous resistant F/F), demonstrating intense levels of pyrethroid resistance in the country (Fotakis et al. 2017). In contrast, the same study observed frequencies of G119S and F290V *ace-1* amino acid changes in 14% and <1% of the mosquitoes, respectively, representing very low resistance to organophosphates and carbamates. In this study, we report a heterozygous and homozygous resistance frequency of 13-44% for *kdr* and 25-50% for AChE insensitivity in *Culex pipiens* mosquitoes from Belgium. It is evident that the rate of resistance is location-dependent and can arise rapidly in mosquito populations given the right selection pressures. In areas with competent vectors and known or suspected arbovirus circulation, it is important to distinguish the types and degrees of insecticide resistance to inform evidence-based decision-making regarding the use of vector control. In most European countries, however, the insecticide resistance profile of *Culex* mosquitoes is an important knowledge gap and potential barrier to effective vector control.

No reports on insecticide resistance in *Culex modestus and Culex torrentium* mosquitoes were found in literature. Interestingly, our study observed a frequency of 43% *kdr* resistant heterozygosity (L/F) in *Culex modestus*, whereas the frequency of G119S *ace-1* was low (3.9% heterozygous). In contrast, we did not observe any resistance alleles in *Culex torrentium* mosquitoes. A larger sample size will be needed to confirm these results, although *Culex torrentium* is not an abundant species found in these locations.

The mutation for *ace-1* G119S was observed at a higher frequency in the heterozygous state compared to the homozygous state. Only one female with the S/S resistant homozygote genotype was found, similar to the findings from Greece (Kioulos et al. 2014). We assume that the high frequency of the *ace-1* G119S heterozygous resistant genotype in *Culex pipiens* is due to the high fitness costs associated when the mutation is present in the homozygote form. This theory is also suggested by other studies where mosquitoes with the S/S homozygote genotype were rarely reported (Weetman et al. 2015). Moreover, under laboratory conditions, a significantly higher mortality was observed in *Anopheles gambiae* pupae with the S/S (homozygote) substitution compared to pupae with the G/G (susceptible) genotype (Djogbénou et al. 2010). As these fitness costs are alleviated in the heterozygote form, repeated selection may lead to a permanent state of heterozygosity in *Culex* populations (Berticat et al. 2002). Another mutation leading to the F290V amino acid substitution was reported to be associated to AChE insensitivity as well, warranting further investigation in Belgian *Culex* mosquitoes (Alout et al. 2007).

A limitation to our study was the small sample size which did not allow us to perform insecticide bioassays such as the WHO cylinder test or CDC bottle bioassay. The detection of these two resistance genotypes does not represent all potential resistance mechanisms that may be involved, such as the changement of the cuticular penetration of insecticides, as well as the degree of phenotypic resistance in these mosquitoes (Donnelly et al. 2009). However, as there is evidence of a correlation between *kdr* resistance genotypes and phenotypic resistance to pyrethroids in *Culex pipiens pipiens*, we expect that there must be some degree of phenotypic resistance to pyrethroids in Belgian *Culex pipiens* with *kdr* L/L or L/F alleles (Tmimi et al. 2018). In comparison with the *Culex pipiens* from Morocco (with 42% frequency of resistant alleles, showing resistance levels 33 times higher than the susceptible strain to permethrin), the *Culex pipiens* from Belgium (with approximately 20% frequency of resistant alleles) are expected to have a lower resistance level to permethrin than the mosquitoes from Morocco but higher than the susceptible mosquitoes from Belgium (Tmimi et al. 2018).

Importantly, the co-occurrence of *kdr* L1014F and *ace-1* G119S suggests that multi-or cross-resistant mosquitoes could be developing in our study area, which may cause an inefficiency of pyrethroid and organophosphate insecticides if used for mosquito control in Belgium. Further measurement of the metabolic activity of enzymes (such as cytochrome p450, esterase and acetylcholinesterase) involved in insecticide detoxification via transcriptomic (Silva Martins et al. 2019) or proteomic methods is warranted to better understand the extent of mosquito insecticide resistance in Belgium (Epelboin et al. 2021).

In the Flanders region of Belgium, a total of 2.4 tons of pesticides were reported to be used in 2020 (Vlaamse Milieumaatschappij 2020). The majority of pesticide use (98%) consisted of herbicides for plant control, of which glyphosate was the most common active ingredient (67%). This was followed by 2,4-dichlorphenoxyacetic acid (2,4-D; 17%), triclopyr (7%), 2-methyl-4-chlorophenoxyacetic acid (MCPA; 4%), and *Bacillus thuringiensis* (2%). The strong use of glyphosate, an organophosphate, may explain the development of *ace-1* G119S resistance in the *Culex pipiens* observed in our study. It was shown that at field-realistic doses, the presence of glyphosate can impair the learning ability of *Aedes aegypti* larvae (Baglan et al. 2018) as well as modify the life history traits and increase insecticide resistance in *Anopheles arabiensis* (Oliver and Brooke 2018). In addition, it was shown that 2,4-D and MCPA can affect AChE sensitivity in humans (Bukowska and Hutnik 2006); therefore, the development of *ace-1* resistance in Belgian *Culex* mosquitoes may also be due to the selection pressures caused by 2,4-D and MCPA. The use of insecticides for insect control in Belgium represented 2% of all pesticide use, specifically for the control of wasps and oak processionary caterpillars. The use of insecticides against mosquitoes was reported in only one location at the French border to prevent the entry of *Aedes* mosquitoes. The most common site for the application of pesticides was open pavement (78%), followed by woody vegetation (13%), and sports fields (5%). In Belgian botanical gardens, the use of fungicides, acaricides and rare pesticides were also reported but not specified. Aside from chemical pesticides, there is evidence that natural plant chemicals and microbes can confer insecticide resistance in mosquitoes (David et al. 2006, Kikuchi et al. 2012). A common botanical pesticide is pyrethrum which, although derived from natural sources, shares a similar structure and mode of action with synthetic pyrethroid insecticides (Isman 2008). Other potential environmental contributors to resistance include natural xenobiotics (allelochemicals), industry pollutants, and domestic insecticides. However, the extent to which Belgian mosquitoes in their adult or immature developmental stages may be exposed to natural pesticides is unclear.

There is little evidence on the efficacy and cost-effectiveness of vector control interventions in Europe (Bellini et al. 2014). Over recent decades, the European Union has been gradually banning the use of insecticides for agriculture, including the pyrethroid permethrin and the organophosphate malathion. In Flanders, there has been a consistent decline in the overall use of pesticides. Since 2010, the quantity of active ingredients has fallen from 15.7 to 2.4 tons in 2020 (Vlaamse Milieumaatschappij 2020). It is possible that a continuous decline of pesticide use may reduce the selection pressure on mosquitoes to develop insecticide resistance. However, the limited use of a small number of active ingredients and the high reliance on glyphosate may facilitate the development of insecticide resistance in mosquitoes if they are being exposed regularly at low concentrations (Oliver and Brooke 2018). As we do not expect the frequency of insecticide resistance in these mosquito populations to decline, we recommend that insecticide resistance levels of mosquitoes are monitored at a larger scale in Belgium and in neighbouring countries as well.

## Conclusion

In this study, we present the first report of *kdr* L1014F and *ace-1* G119S insecticide resistance in *Culex pipiens, Culex modestus*, and *Culex torrentium* mosquitoes from Belgium. We highlight the importance of mosquito surveillance for the development and implementation of effective arbovirus control strategies in the event of an outbreak.

## Acknowledgements

This project was funded by KU Leuven (C14/20/108 and starting grant STG/19/008). We would like to thank Prof. Johan Neyts (Rega Institute, KU Leuven) for allowing us the use of his laboratory space and equipment for our experimental work. We would like to thank the City Green Management of Leuven and KU Leuven for providing the permits for the mosquito field collections from 2019 to 2021.

